# Controlling emulsification for organic solvent-based tissue clearing

**DOI:** 10.1101/628925

**Authors:** Bing Hou, Xinyi Liu, Dan Zhang, Zaifu Yang, Hui Hui, Peixin Wang, Jiarui Wang, Zilong Qiu, Yanyan Liu, Zhengyi Yang, Tianzi Jiang

**Affiliations:** Brainnetome Center, Institute of Automation, Chinese Academy of Sciences, Beijing 100190, China; National Laboratory of Pattern Recognition, Institute of Automation, Chinese Academy of Sciences, Beijing 100190, China; Beijing Institute of Radiation Medicine, Beijing 100850, China; Key Laboratory of Molecular Imaging, Chinese Academy of Sciences, Beijing, 100190, China; Department of Neurosurgery, PLA General Hospital, Beijing 100853, China; Department of Neurosurgery, PLA 306^th^ Hospital, Beijing 100853, China; Institute of Neuroscience, Chinese Academy of Sciences, China; Queensland Brain Institute, The University of Queensland, Brisbane, QLD 4072, Australia; The Clinical Hospital of Chengdu Brain Science Institute, MOE Key Lab for Neuroinformation, University of Electronic Science and Technology of China, Chengdu 625014, China; CAS Center for Excellence in Brain Science and Intelligence Technology, Chinese Academy of Sciences

## Abstract

Organic solvent-based tissue clearing techniques are hampered by the quenching of fluorescent proteins, partially due to routine complete dehydration. Unexpectedly, we discovered that complete dehydration is unnecessary for organic solvents to clear tissues and that the hidden purpose has been to prevent emulsification. After controlling emulsification of organic solvent-cleared but incompletely dehydrated mouse brain, we achieved sufficient tissue transparency that allowed light-sheet imaging while well-preserving the fluorescence of fluorescent proteins.

## Manuscript

Tissue optical clearing techniques are powerful tools for investigating the inherent three-dimensional structure of cells and organs^1^. Compared with water-based counterparts, organic solvent-based clearing techniques can render tissue extremely transparent, which is particularly suited to rapid three-dimensional imaging, such as light-sheet microscopy^2–4^. However, the fact that organic solvent-based clearing techniques quench fluorescent proteins makes them less useful, since a substantial number of biological materials under investigation have been labeled with fluorescent proteins.

To overcome this limitation, a straightforward approach is to screen for new dehydrants and clearing solvents, such as tetrahydrofuran (THF) and dibenzyl ether (DBE), that have a weaker quenching effect on fluorescent proteins ^3,5^. However, THF and DBE cannot stably preserve the endogenous fluorescence of fluorescent proteins; thus the fluorescence imaging of fluorescent proteins in cleared tissues remains problematic^6–8^. Another promising attempt is to add suitable organic bases^9^ or organic antioxidants^10^ to solvents to retain fluorescent proteins in the fluorescent state. This strategy has recently been adopted to help fluorescent proteins in organic solvent-cleared tissues maintain their fluorescence over weeks to months^9,10^.

An alternative approach is to replace organic solvent-susceptible fluorescent proteins with organic solvent-resistant synthetic fluorophors. Immunostaining using fluorophor-labeled antibodies can bypass the quenching problem associated with fluorescent proteins and enable stable fluorescence imaging of cleared tissues ^4,11,12^. Although this modification has the advantage of allowing the researcher to identify numerous molecular targets, it is hardly applicable for large volume adult mammalian organs due to the low level of antibody penetration ^4^.

To explore a different strategy, we investigated whether complete dehydration, one of the main reasons for the quenching of fluorescent proteins ^6,13,14^, was indispensible for organic solvent-based tissue clearing techniques. Typically, organic solvent-based clearing techniques comprise two stages: dehydration with an ascending gradient of amphiphilic dehydrants and transparentization with water-insoluble organic solvents with a high refractive index. To achieve excellent tissue transparency, previous reports consistently suggested, but without explanation, that dehydration should be conducted as completely as possible for organic solvent-based tissue clearing techniques ^2–5,15^, implying that there might be a key but unidentified mechanism accounting for the pursuit of complete dehydration.

To investigate whether and how remaining water influences tissue transparency, we dehydrated an adult mouse brain through the THF gradient following the 3DISCO protocol, as was done previously ^3^, except that we ended at 97% THF (vol / vol) and continuously observed its transparency after it was immersed in DBE. Unexpectedly, almost all parts of the brain parenchyma, including the deep subcortical structures, were rendered transparent within 45 minutes but thereafter turned opaque again (**Supplementary Fig. 1**). Tissue opacity is known to result mainly from light scattering due to the inhomogeneity of water, protein, lipid, or other scatters in the tissue ^1^. However, none of the known mechanisms could readily explain the re-opacification.

Based on the properties (**Supplementary Table 2**) and uses of chemical reagents involved in organic solvent-based tissue clearing techniques, we hypothesized that the re-opacification might arise from emulsification, which refers to a uniform blend of two or more liquids that are normally immiscible. In other words, emulsification could develop inside organic solvent-cleared biological samples if complete dehydration could not be guaranteed.

To verify our hypothesis, we performed a microscopic examination of brain slices and revealed numerous water droplets inside the re-opacified tissues but not inside the transparent tissues (**Supplementary Fig. 2**), indicating emulsification formation during re-opacification. We further conducted simulation tests in the tubes to assess whether emulsification could develop when a tiny amount of water was directly mixed with clearing solvents. Since DBE was recommended for organic solvent-based clearing techniques ^3,4^, we chose this established clearing agent to start our test. We found without exception that emulsification happened at 23°C immediately after water was fully mixed with DBE in a given volume ratio of 1:50 with fierce shaking, irrespective of the presence of tested dehydrants such as THF (**Fig. 1a**). The resultant emulsion contained numerous fine water droplets with a diameter of 2-4 μm (**Supplementary Fig. 3**), which caused scattering and rendered the emulsion milky. When the emulsion was heated mildly, its milky appearance faded away as the temperature rose to 35°C (**Fig. 1b-c**). This is indicative of the temperature-sensitive nature of emulsification formation.

**Figure 1.**
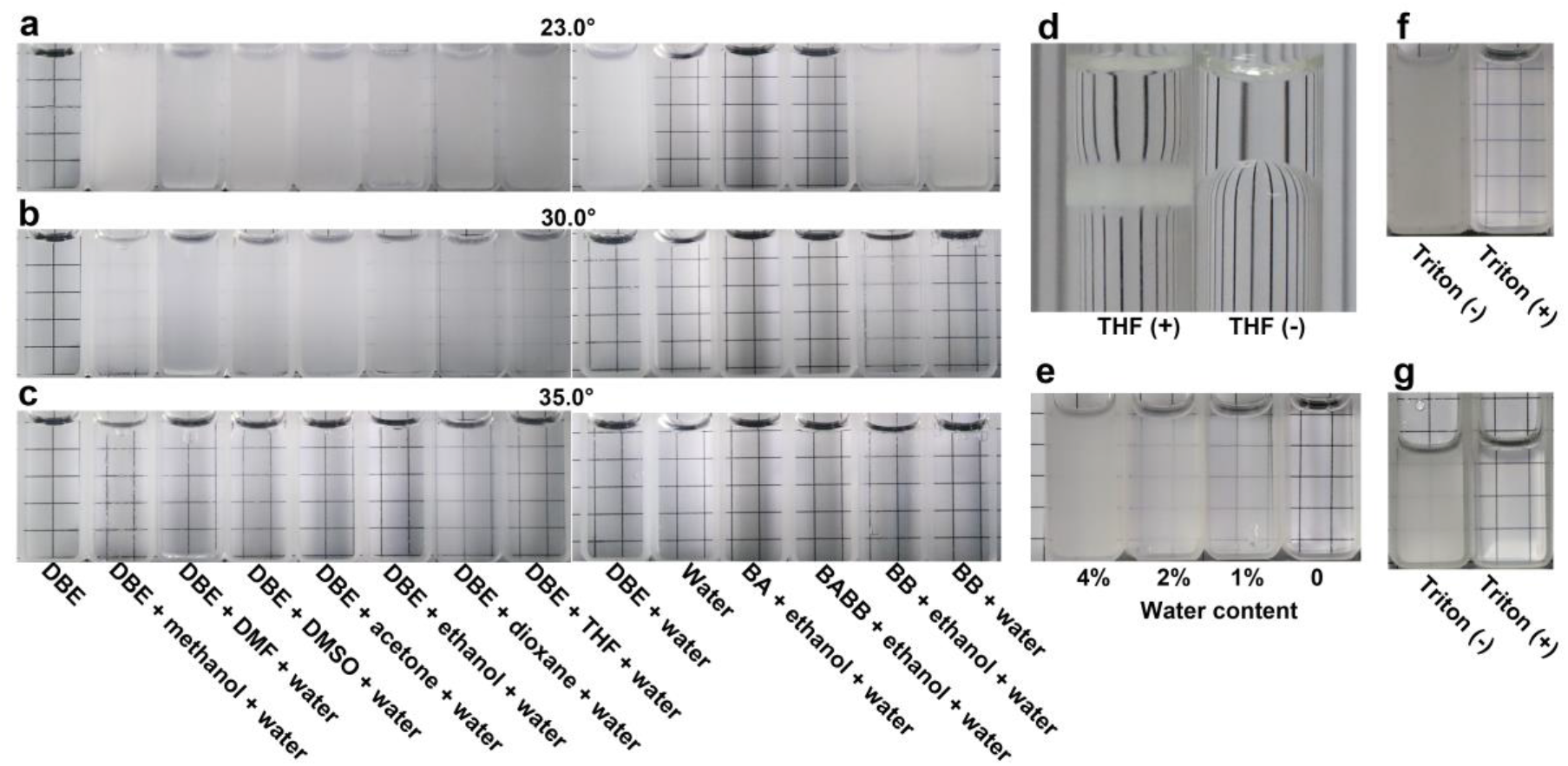
Mixture of clearing agents and water leading to emulsification. (a-c) The solubility of water in the agents influenced the extent of emulsification in a temperature-dependent manner when water was completely mixed with different clearing agents. At 23°C, emulsification happened immediately after water was fully mixed with DBE or BB in a given volume ratio of 1:50, with or without any dehydrant tested but did not happen after water was fully mixed with BA or BABB (a). The milky appearance differentially faded in different mixtures at 30°C in relationship to the solubility of water in the different clearing agents (b) and without fail disappeared at 35°C (c). (d) Emulsification immediately occurred at the interface when water (above) containing THF was gently added onto DBE (below), resulting in a sharp partition split between pure water (above) and DBE (below) without emulsification formation. Note that the vertical line behind the glass tube is deformed due to the different refractive indices of water (1.33) and DBE (1.56). (e) The severity of emulsification is dependent on the amount of water in the DBE. (f) Emulsification occurred when water alone was fully mixed with DBE but did not occur when water containing Triton X-100 was fully mixed with DBE. (g) Emulsion resulting from a well-mixed mixture of water and DBE was rendered transparent after addition of Triton X-100.

Before DBE was discovered to be a clearing solvent, BABB, a cocktail of benzyl alcohol (BA) and benzyl benzoate (BB), had been selected to clear biological samples ^2^. In the same 1:50 ratio as above, a well-mixed mixture of water with BB also emulsified, but a well-mixed mixture of water with BA or BABB did not (**Fig. 1a**). Although clearing agents such as BA and DBE are nominally called “water-insoluble” solvents, they can indeed dissolve a certain amount of water depending on differences in its solubility in these agents (**Supplementary Table 2**). Thus, organic solvent-based tissue clearing would be at risk of emulsifying if the content of water remaining in the tissues after dehydration exceeded water’s solubility in that specific clearing agent.

Full dispersion is required for two mutually immiscible liquids to induce emulsification, but mechanical approaches to induce full dispersion, such as fierce shaking, were actually absent during the realistic organic solvent-based tissue clearing. Noting their amphiphilic nature, we inferred that the dehydrants used might also act as emulsifiers that remarkably facilitated the full dispersion of water into clearing agents (**Fig. 1b**). This inference was soon supported when emulsification immediately happened after a well-mixed mixture of THF with water was gently added onto DBE but did not when water alone was gently added (**Fig. 1d**).

We then found that the severity of emulsification depended on the existing water content in the clearing solvents. In the absence of water, a well-mixed mixture of DBE with THF did not induce emulsification (**Fig. 1e**), suggesting that complete dehydration should be effective for preventing emulsification in organic solvent-cleared organs. In addition to complete dehydration, enhancing water’s solubility in the clearing agents by heating also reduced emulsification (**Fig. 1b-c**). Moreover, *ex ante* or *ex post* addition of Triton X-100, one of the surfactants capable of blocking water molecule aggregation in organic solvents, could also prevent or eliminate emulsification and render the emulsion transparent (**Fig. 1f-g**).

We then examined whether these strategies aimed at preventing emulsification or demulsification could also secure tissue transparency in realistic organic solvent-based tissue clearing. We found that the whole adult rat or mouse brains did not emulsify even at a lower temperature (18°C) when complete dehydration was finished after three rounds of 100% THF following the standard 3DISCO procedure ^3^ (**Fig. 2a, d**). However, when the brain samples were dehydrated after only one round of 100% THF, they were transparent at 26°C but turned less transparent at 18°C due to extensive but slight emulsification (**Fig. 2b**). In accordance with the results in the tube tests, mild heating (35°C) eliminated emulsification in the samples and rendered the samples highly transparent again (**Fig. 2b**). Moreover, adding Triton X-100 to the clearing agent also improved tissue transparency (**Fig. 2c**).

**Figure 2.**
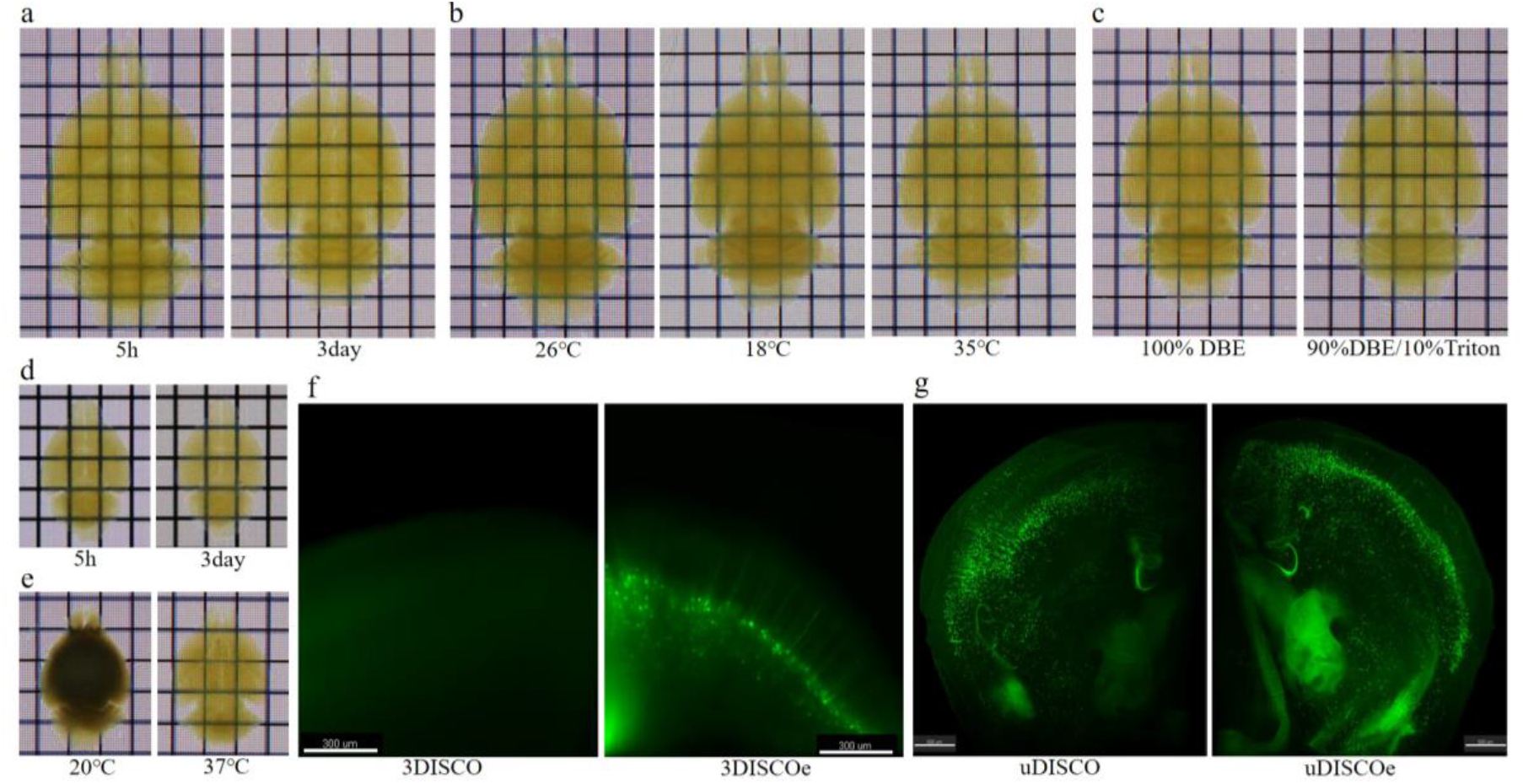
Emulsification control in organic solvent-based tissue clearing. (a) The whole adult rat brains cleared by standard 3DISCO remained transparent without emulsification formation for 3 days even at a lower temperature (18°C) when complete dehydration was finished after three rounds of 24-hour incubation in 100% THF. (b) When the rat brain samples were dehydrated through only one round of 100% THF, they were rendered transparent at 26°C but turned less transparent at 18°C due to extensive but slight emulsification and then became highly transparent again after mild heating (35°C). (c) Addition of Triton X-100 in the clearing agent also improved tissue transparency of the rat brain dehydrated through one round of 99% THF. (d) The whole mouse brains cleared by standard 3DISCO remained transparent without emulsification formation for 3 days when complete dehydration was finished after three rounds of 12-hour incubation in 100% THF. (e) Our method rendered the whole adult mouse brain transparent within 6 hours (right), as compared with 3-4 days for the original protocol (left). (f) The fluorescence signal of green fluorescent proteins was bright in the DBE-cleared brains after final dehydration with 99% THF, whereas no fluorescence signal was detected in the cleared brains after complete dehydration. (g) Incomplete dehydration also markedly enhanced the fluorescence signal of green fluorescent proteins in the brain slices that were cleared using a cocktail of BABB and diphenyl ether (DPE).

We revisited the experimental results shown in Fig. S1 and were able to explain the entire process of clearing and re-opacification. The clearing started from the surface of the brain (olfactory bulbs, cerebellum and outer layers of the cerebral cortex) and extended gradually to the subcortical nuclei. The re-opacification followed the same pattern. We could see that when the deep brain structures cleared at 45 min, the transparency of the olfactory bulbs and cerebellum started to decrease. The spatiotemporal pattern of the transparency of the brain could be explained as the dehydrant diffusing into the tissue to clear it, followed by the dehydrant causing emulsification to opacify it. One may miss the clearing status, which all the brain structures underwent but at different times, before re-opacification, if the process is not observed continuously. This could lead to the misunderstanding that complete dehydration is a prerequisite condition for tissue clearing.

These findings from tissue experiments appear to have confirmed our inference that preventing emulsification was indeed the real reason for pursuing complete dehydration during organic solvent-based tissue clearing. This proven mechanism could allow researchers to tailor the procedure of organic solvent-based clearing techniques on demand. For example, to speed up tissue clearing, investigators could choose to replace complete dehydration, the most time-consuming stage during the whole procedure, with incomplete dehydration plus mild heating and surfactant addition. As a result of these investigations, we found that the new protocol could render the whole adult mouse brain transparent within only 6 hours (**Fig. 2e**) compared with 3-4 days for the original protocol^3^ (**Fig. 2d**).

In contrast to the popular belief that complete dehydration is necessary for organic solvents to achieve satisfactory tissue transparency ^2–5,15^, our experiments demonstrated that samples that were cleared by organic solvents instead allowed the presence of a tiny amount of water. This unexpected finding inspired us to utilize the remaining water to protect fluorescent proteins from quenching ^6,13,14^. Thereafter, we examined whether retaining a tiny amount of water would enable fast and stable fluorescence imaging for large volume mammalian tissues that were cleared by organic solvents.

To clear a given organ, its transparency should meet the minimal requirement for the desirable imaging method. As compared with confocal and two-photon microscopy, light-sheet microscopy (LSM) allows at least 100 times faster imaging speed but requires higher tissue transparency ^16^. In view of LSM’s superiority in whole organ imaging, we rendered the mouse brain transparent enough for LSM after controlling the extent of emulsification by adjusting the water content in the tissue. We found that the fluorescence signal of fluorescent proteins was bright in the DBE-cleared brains after final dehydration with 99% THF (**Fig. 2f, Supplementary Movie 1**) whereas we did not detect a fluorescence signal in the cleared brains after complete dehydration. Moreover, incomplete dehydration also remarkably enhanced the fluorescence signal of fluorescent proteins in the brain slices that were cleared using the recently established protocol that utilizes a cocktail of BABB and diphenyl ether (DPE) ^10^ (**Fig. 2g**). This finding suggests that controlling emulsification is an effective and general strategy for organic solvent-based tissue clearing techniques while avoiding the quenching of fluorescent proteins.

In conclusion, we present here a simple but robust solution by which organic solvents can achieve efficient tissue clearing while well-preserving endogenous fluorescence signals, a process which had been thought to be intrinsically challenging ^17^. This solution was inspired by our discovery that complete dehydration was not indispensible for organic solvents to transparentize tissues. In fact, the hidden purpose of routine complete dehydration has been to prevent emulsification, which explains why complete dehydration had been empirically suggested for organic solvents to clear tissues. Our work showed that complete dehydration is just one approach to preventing emulsification. Combined with other approaches that aimed to prevent or eliminate emulsification, incomplete dehydration can also render the tissues transparent but consume much less time than complete dehydration. Interestingly, the extent of emulsification can be controlled on demand by adjusting the water content in incompletely dehydrated tissues by considering water’s solubility in the clearing agents used. More importantly, the remaining water can protect fluorescent proteins from quenching. Informed with this knowledge, researchers can create the tissue transparency required for light-sheet imaging of organic solvent-cleared organs while the fluorescent signal of fluorescent proteins is kept bright and stable.

## Methods

Methods and any associated references are available in the online version of the paper.

Note: Any Supplementary Information and Source Data files are available in the online version of the paper.

## Acknowledgments

The authors thank Profs. Shan Yu and Bing Liu for the insightful discussion, Prof. Yousheng Shu for the use of the laboratory and Rhoda E. and Edmund F. Perozzi, PhDs for editing the manuscript. This work was partially supported by the Natural Science Foundation of China (Grant Nos. 31620103905, 91732305, and 81671855), the Science Frontier Program of the Chinese Academy of Sciences (Grant No. QYZDJ-SSW-SMC019), National Key R&D Program of China (Grant No. 2017YFA0105203).

## Author Contributions

B.H. proposed the concept and drafted the manuscript.

B.H., D.Z. and Z.Y. designed the experiments.

D.Z., P.W., J.W., Y. L., and X.L. prepared the biological samples.

B.H., D.Z., Z.Y. and J.W. performed the tube and tissue experiments.

X.L., D.Z. and H.H. performed the light-sheet imaging.

Z.Y. made critical revisions in the preparation of the manuscript.

T.J. designed the concept and protocols, supervised the project and experiments and wrote the manuscript.

## Competing financial interests

The authors (B.H., D.Z., Z.Y. and T.J.) have applied for patents on the emulsification control methods for organic solvent-based tissue clearing, assigned to the Institute of Automation, Chinese Academy of Sciences.

## Online Methods

### Reagents and reagent preparation

Reagents, if not otherwise indicated, were purchased from Aladdin Industrial Corporation (Shanghai, China), Alfa Aesar Chemicals Company (Shanghai, China), or Sinopharm Chemical Reagent Company (Beijing, China). The detailed information about the CAS number, supplier, and catalog number of these reagents is listed in **Supplementary Table 1**. The physical and chemical properties of these reagents are shown in **Supplementary Table 2**. For safety’s sake, experimenters should use proper gloves and mask when handling THF or other organic solvents, avoiding inhalation or contact with skin and eyes. The organic solvents must be prepared in a fume hood and stored in a glass bottle with a compatible lid. Before use, peroxides in THF were removed by column absorption chromatography with basic aluminum oxide so as to be undetectable by Quantofix Peroxide 25 test sticks (Z249254, Sigma-Aldrich, Germany) whose minimum detection level for peroxides was 0.5 mg/L.

### Animals

All animals were housed and treated in accordance with institutional guidelines. The experimental procedures and housing conditions were approved by the Animal Experiment Committee of the local authority. Wild-type (C57BL/6N) and Thy1-YFP (line H) mice in a C57BL/6N background were used for optical clearing. Wild-type (Sprague-Dawley) rats were also used. Adult mice or rats over 70 days of age were deeply anesthetized with an intraperitoneal overdose of sodium pentobarbital (150 mg/kg body weight) and transcardially perfused with 1×PBS followed by 4% (wt/vol) paraformaldehyde (PFA) in 1×PBS. The whole brains were excised and then post-fixed in the same fixative at 4 °C overnight.

### Tube tests for investigating emulsification

A series of experiments were carried out to reveal the effect of the residual water on tissue transparency and the mechanism of emulsification control.

First, to determine whether the clearing agent and water can be emulsified, we placed the liquid mixed according to the formulations in Supplementary Table 3 in a glass tube having a capacity of 20 ml and shook it well. The liquid mixture was drawn and transferred to a dry, clean quartz beaker to observe its transparency. We recorded the temporal change in transparency of different liquid mixtures at different temperatures, to see whether the emulsification is temperature sensitive.

Second, to elucidate the effects of the dehydrant, water content, and surfactant on emulsion formation, we carefully placed the liquids, separately mixed according to the formulations in Supplementary Table 4, 5, and 6 in glass test tubes with a capacity of 20 ml. The liquids that were mixed with water were thoroughly oscillated. The liquid mixture was drawn and transferred to a dry, clean quartz beaker to observe the transparency of the mixture. The change in transparency of the interface between the different liquids was observed.

Third, to look for the presence of water droplet formation, a small amount of the mixed solution was taken and observed under a microscope. To look for the presence of water droplet formation in the transparent and re-opaque samples, the animal brain was cut with a blade into tissue slices about 2 mm thick, dehydrated according to the procedure of 3DISCO with emulsification control, and observed under a microscope.

### Optical tissue clearing using 3DISCO and its variants

Using the standard 3DISCO^3^ protocol, PFA-fixed whole mouse brain samples were serially immersed in 30-50 ml of 50%, 70%, 80%, and 95% (vol/vol) THF, each for about 12 h with gentle rotation at 26 °C. The whole mouse brain samples were then immersed in 100% (vol/vol) THF for 12 h for 3 rounds and finally in DBE until they were transparent. For the whole rat brains, each step was extended to 24 hours.

To achieve fast 3DISCO clearing, a hemisphere sample of Thy-1-YFP(H line) mouse brain was sequentially dehydrated by 50%THF, 70%THF, 80%THF, 95%THF, and 99%(vol/vol)THF, about 90 min per step at 20°C. The sample was scanned by light-sheet microscopy after 30 min immersion in DBE.

In the 3DISCO variant with controlled emulsification (3DISCOe for short), the brain sample was soaked in an ascending gradient dehydrate solution - 50%THF, 70%THF, 80%THF, 95%THF, 97%THF - for about 12 h per step. The dehydration ended at 97%THF rather than 100% THF. Furthermore, the sample was transferred into DBE in a glass dish 15 cm in diameter. Paper with a grid of 5×5 mm was put under the glass dish for observing volumetric and transparency changes. The photographic recording of the changes started immediately after a whole mouse brain was immersed in DBE. Pictures were taken every 5 min.

### Optical tissue clearing using uDISCO and its variants

uDISCO was performed following the original protocol. A brain sample of Thy-1-YFP (H line) mouse was immersed in 30%, 50%, 70%, 80%, 90%, 96%, and 100% (vol/vol) TBA for 12 h at 34°C in sequence. After one hour degreasing in DCM, the sample was transferred into BABB-D4 at room temperature. In uDISCO with emulsification control (uDISCOe for short), 100% TBA was omitted as complete dehydration was not essential for tissue clearing. The other conditions were the same as the standard uDISCO. The samples were observed until complete transparency. Foil paper was used to protect the samples from light throughout the entire experiment.

### Imaging with light-sheet microscopy

The cleared brains were fixed on sample holders by a screw and immersed in tissue clearing solutions held by reservoir. The transparent tissues were imaged using a light-sheet fluorescence microscope (UltraMicroscope II, LaVision BioTech) equipped with a binocular body (Olympus, MVX10), a 2x objective (Olympus, NA = 0.5, working distance = 5.7 mm), and a lens cap (LaVision Biotech). In the experiments, an excitation laser with a wavelength of 470 nm was selected for the scanning. Imaris 9.0.1 and ImageJ were used for image analysis and presentation.

## Supplementary Information

**Supplementary Figure 1.**
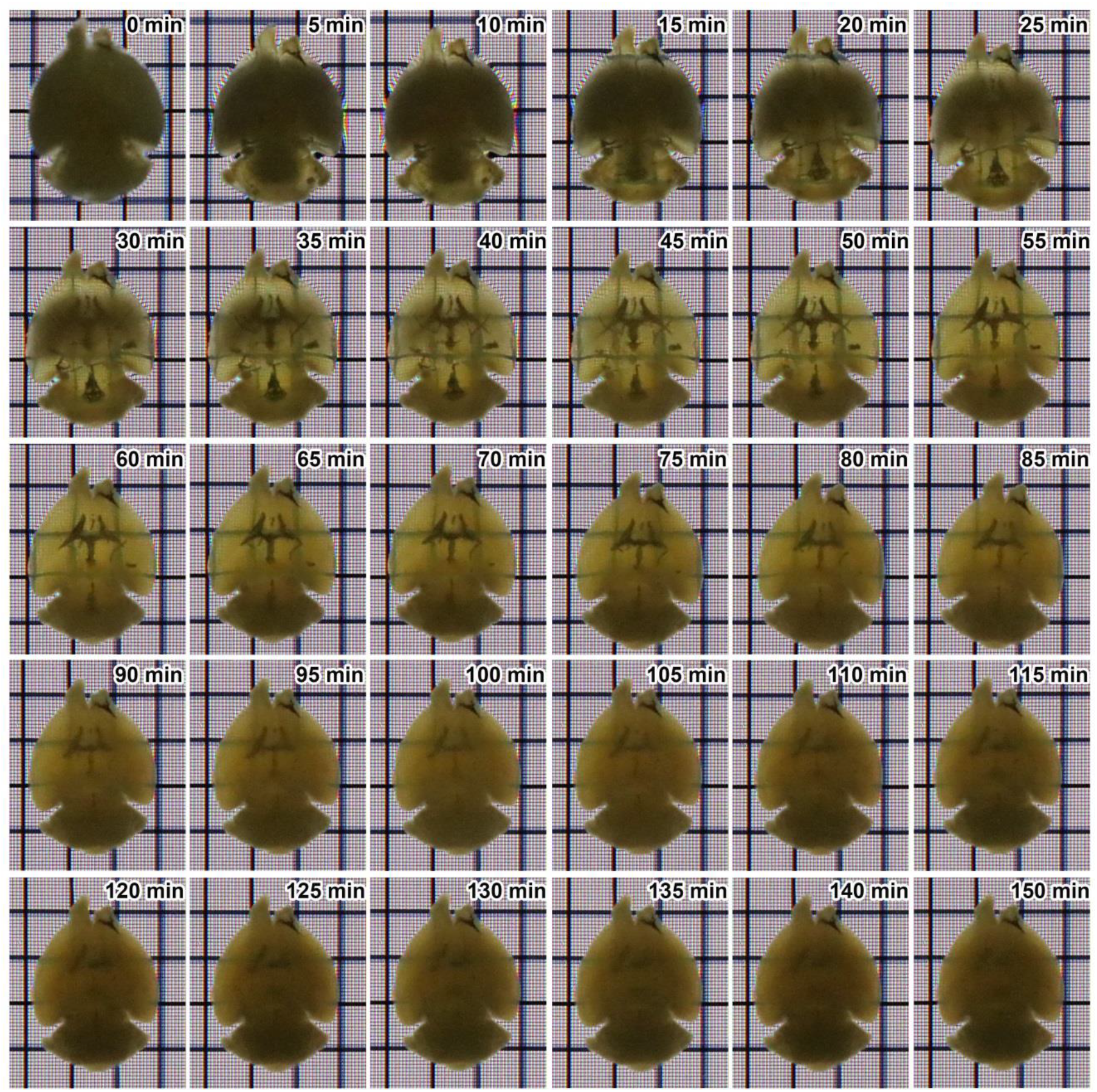
Dynamic changes in tissue transparency for an incompletely dehydrated mouse brain. An adult mouse brain was incompletely dehydrated through an ascending THF gradient ending at 97% THF (vol / vol) and then immersed in DBE. Unexpectedly, almost all parts of the brain parenchyma, including deep subcortical structures such as the striatum, could be rendered transparent within 45 minutes but thereafter turned opaque again. Note that the brain ventricles remained opaque due to the presence of cerebrospinal fluid rich in water. Grid size: 5× 5 mm.

**Supplementary Figure 2.**
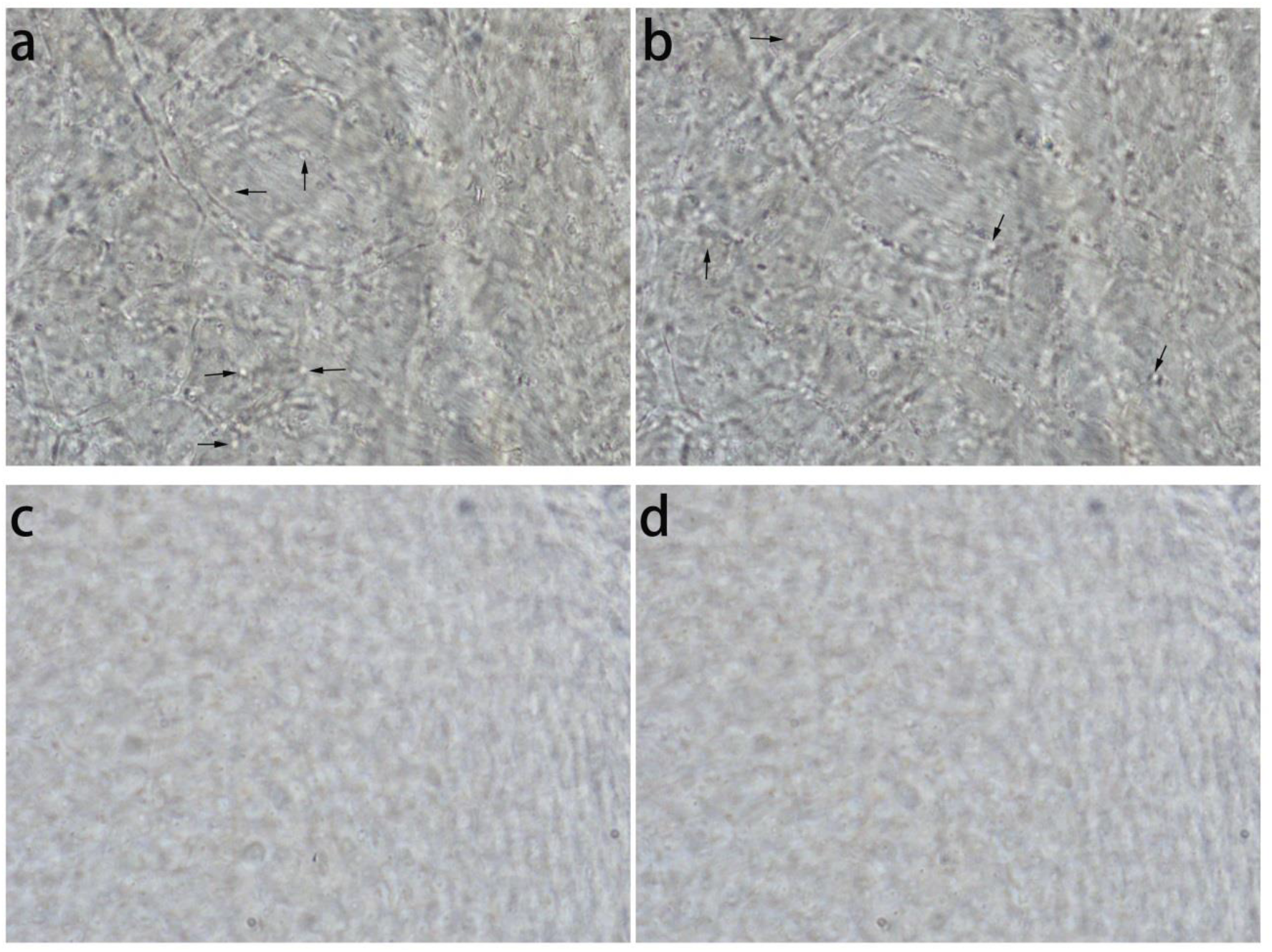
Microscopic examination of brain slices. Microscopic examination of brain slices revealed numerous water droplets (black arrows) inside the re-opacified tissues (a, b) but not inside the transparent tissues (c, d). Interval between the upper two images and the lower two images were both 9 μm in depth.

**Supplementary Figure 3.**
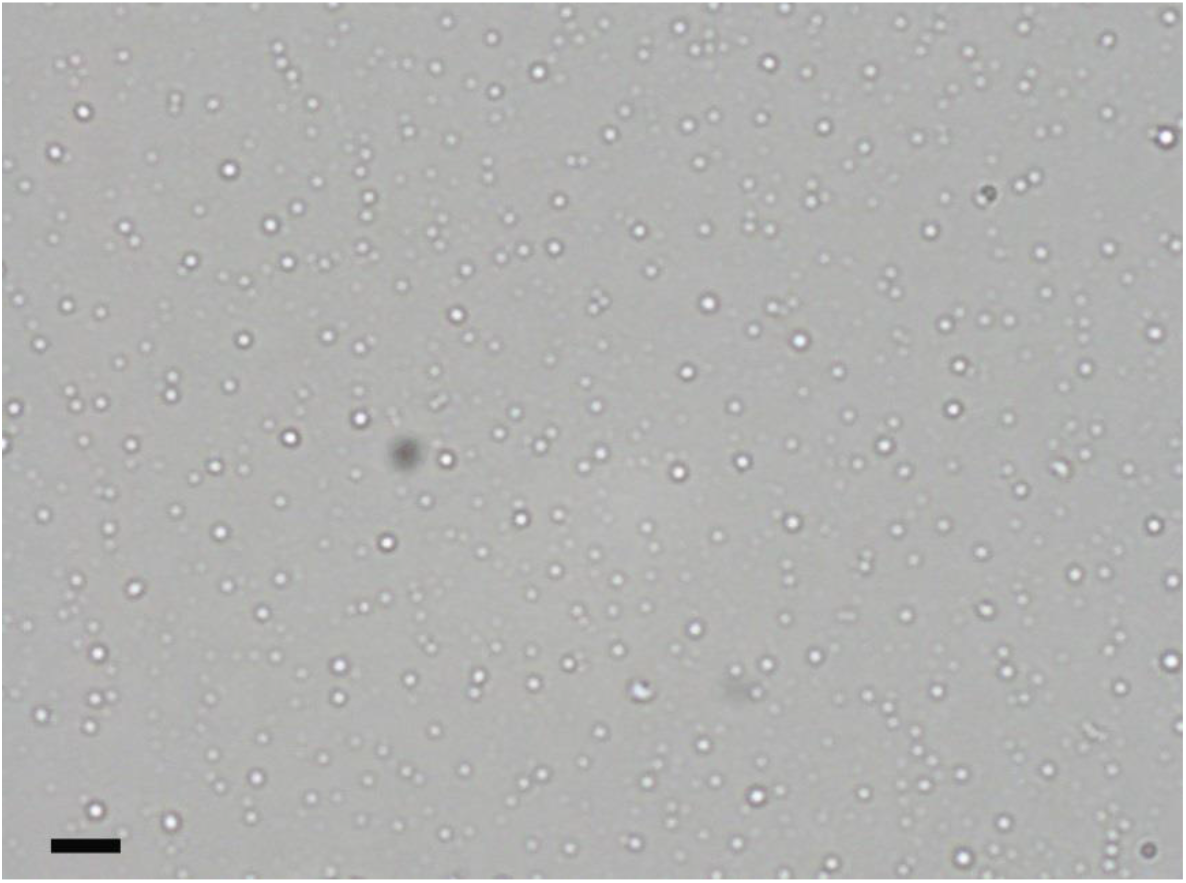
Microscopic examination of emulsion. Numerous water droplets (white dots) with a diameter of several μm were recognized under microscopic examination in emulsion resulting from a full mixture of water, THF, and DBE. Scale bar: 10 μm.

**Supplementary Table 1.** List of chemical reagents used

| **Reagent (abbreviation)** | **CAS number** | **Supplier** | **Catalog number** |
| --- | --- | --- | --- |
| Acetone | 67-64-1 | Sinopharm Chemical Reagent Co. | 10000492-500ml |
| Aluminum oxide | 1344-28-1 | Aladdin Industrial Corporation | A100282-500g |
| Benzyl alcohol (BA) | 100-51-6 | Sinopharm Chemical Reagent Co. | 30020618-500ml |
| Benzyl benzoate (BB) | 120-51-4 | Sinopharm Chemical Reagent Co. | 30020828-500ml |
| n-Butanol | 71-36-3 | Sinopharm Chemical Reagent Co. | 10005218-500ml |
| Dibenzyl ether (DBE) | 103-50-4 | Alfa Aesar (China) Chemicals Co. | 10186739-500g |
| Dichloromethane (DCM) | 79-09-2 | Sinopharm Chemical Reagent Co. | 80047392-500ml |
| Dimethyl formamide (DMF) | 68-12-2 | Sinopharm Chemical Reagent Co. | 81007718-500ml |
| Dimethyl sulfoxide (DMSO) | 67-68-5 | Sinopharm Chemical Reagent Co. | 30072418-500ml |
| Dioxane | 123-91-1 | Sinopharm Chemical Reagent Co. | 10008992-500ml |
| Diphenyl ether (DPE) | 101-84-8 | Aladdin Industrial Corporation | D110644-500ml |
| Ethanol | 64-17-5 | Sinopharm Chemical Reagent Co. | 10009292-500ml |
| Formamide | 75-12-7 | Sinopharm Chemical Reagent Co. | 30091218-500ml |
| Methanol | 67-56-1 | Sinopharm Chemical Reagent Co. | 10014192-500ml |
| Tert- butyl alcohol | 75-65-0 | Aladdin Industrial Corporation | T110351-500ml |
| Tetrahydrofurane (THF) | 109-99-9 | Aladdin Industrial Corporation | T103263-500ml |
| Triton X-100 | 9002-93-1 | Aladdin Industrial Corporation | T109026-500ml |

**Supplementary Table 2.** Physical and chemical properties of dehydrants and clearing agents

| Reagent | Density (g/ml) | Molecular weight | Refractive index | Polarity | Solubility in water at 20°C |
| --- | --- | --- | --- | --- | --- |
| Water | 1.00 | 18 | 1.33 | 9.0 | -- |
| Formamide | 1.13 | 45 | 1.45 | 7.3 | Miscible |
| Methanol | 0.79 | 32 | 1.33 | 6.6 | Miscible |
| Dimethyl sulfoxide (DMSO) | 1.10 | 78 | 1.48 | 6.5 | Miscible |
| Dimethyl formamide (DMF) | 0.94 | 73 | 1.43 | 6.4 | Miscible |
| Ethanol | 0.79 | 46 | 1.36 | 5.2 | Miscible |
| Acetone | 0.79 | 58 | 1.36 | 5.1 | Miscible |
| Dioxane | 1.03 | 88 | 1.42 | 4.8 | Miscible |
| Tetrahydrofurane (THF) | 0.89 | 72 | 1.41 | 4.2 | Miscible |
| n-Butanol | 0.81 | 74 | 1.40 | 3.9 | 7.7 g/100 ml |
| Dichloromethane (DCM) | 1.33 | 85 | 1.42 | 3.4 | 2.0 g/100 ml |
| Benzyl alcohol (BA) | 1.04 | 108 | 1.54 | NA | 4.0 g/100 ml |
| Benzyl benzoate (BB) | 1.12 | 212 | 1.57 | NA | NA |
| Dibenzyl ether (DBE) | 1.04 | 198 | 1.56 | 3.3 | 0.1 g/100 ml |
| Diphenyl ether (DPE) | 1.08 | 170 | 1.58 | NA | NA |

NA: data not acquired

Data source:

http://www.inchem.org

https://cameochemicals.noaa.gov

http://www.chemicalland21.com

**Supplementary Table 3.**
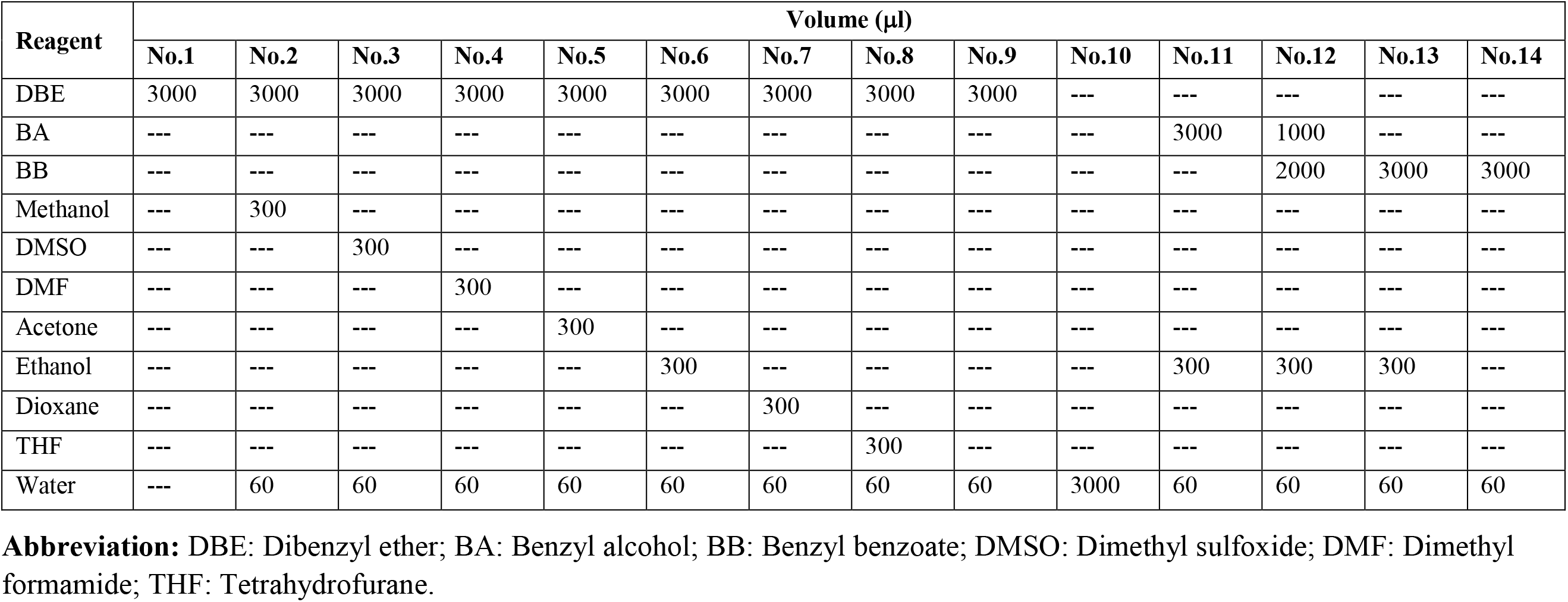
The proportion of reagents in the tubes numbered from left to right used in Figure 1a-c

**Supplementary Table 4.**
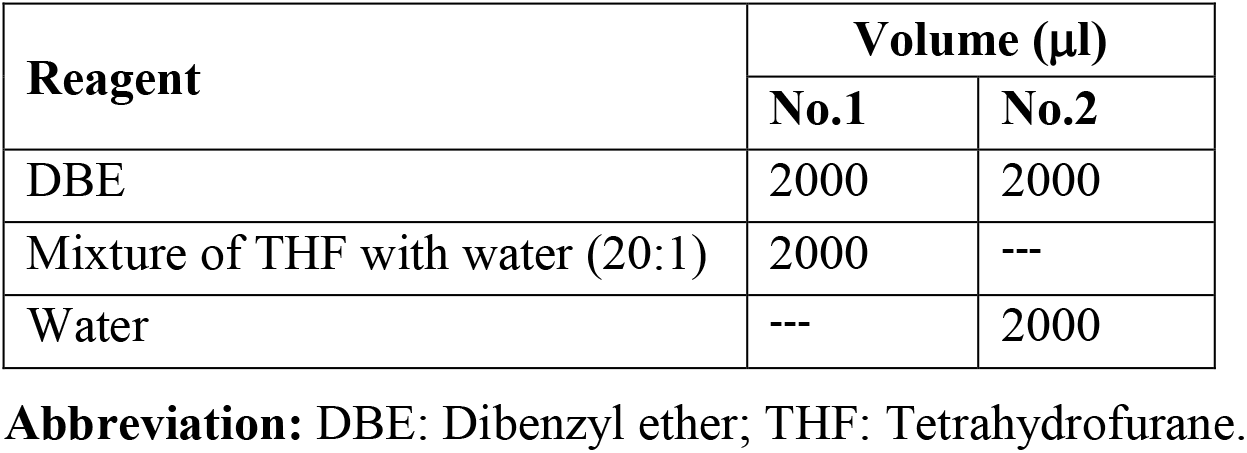
The proportion of reagents in the tubes numbered from left to right in Figure 1d

| Reagent | Volume (μl) |  |
| --- | --- | --- |
|  | No.1 | No.2 |
| DBE | 2000 | 2000 |
| Mixture of THF with water (20:1) | 2000 | --- |
| Water | --- | 2000 |

**Abbreviation:** DBE: Dibenzyl ether; THF: Tetrahydrofurane.

**Supplementary Table 5.** The proportion of reagents in the tubes numbered from left to right in Figure 1e

| Reagent | Volume (μl) |  |  |  |
| --- | --- | --- | --- | --- |
|  | 4% | 2% | 1% | 0 |
| DBE | 3000 | 3000 | 3000 | 3000 |
| THF | 300 | 300 | 300 | 300 |
| Water | 120 | 60 | 30 | 0 |

**Abbreviation:** DBE: Dibenzyl ether; THF: Tetrahydrofurane.

**Supplementary Table 6.** The proportion of reagents in the tubes numbered from left to right in Figure 1f

| Reagent | Volume (μl) |  |
| --- | --- | --- |
|  | No.1 | No.2 |
| DBE | 750 | 750 |
| Mixture of THF with water (20:1) | 150 | 150 |
| Triton X-100 | --- | 150 |

**Abbreviation:** DBE: Dibenzyl ether.

**Supplementary Table 7.** The proportion of reagents in the tubes numbered from left to right in Figure 1g

| Reagent | Volume (μl) |  |
| --- | --- | --- |
|  | No.1 | No.2 |
| DBE | 500 | 500 |
| Emulsion* | 350 | 350 |
| Triton X-100 | --- | 80 |

\*: Emulsion derived from blend of 750 μl DBE and 150 μl mixture of THF with water.

### Supplementary Movies

**Movie S1:**
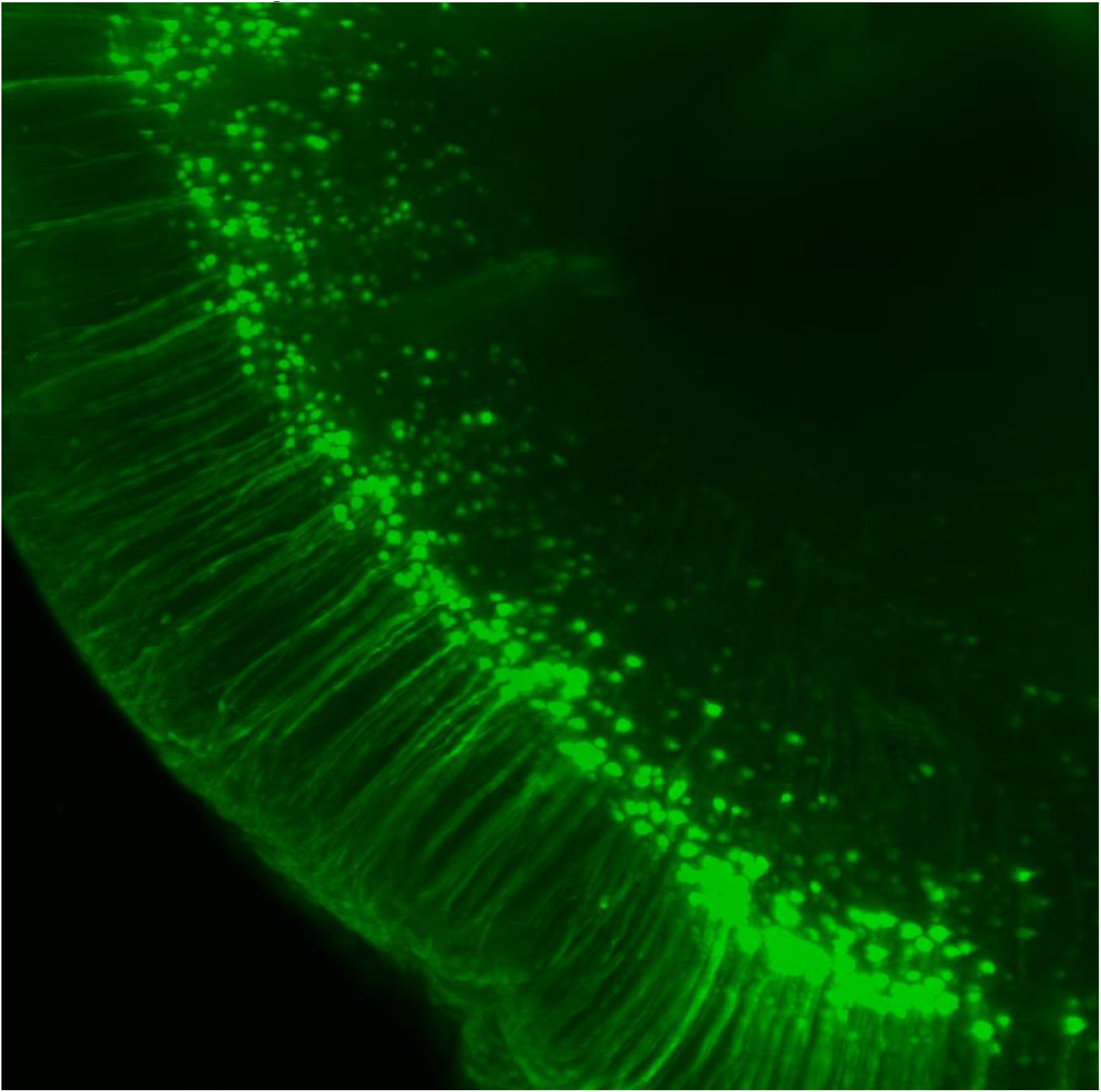
Movie.S1.gif

