## Supplementary figures and images for "Controlling emulsification for organic solvent-based tissue clearing"

### Supplemental Movie 1

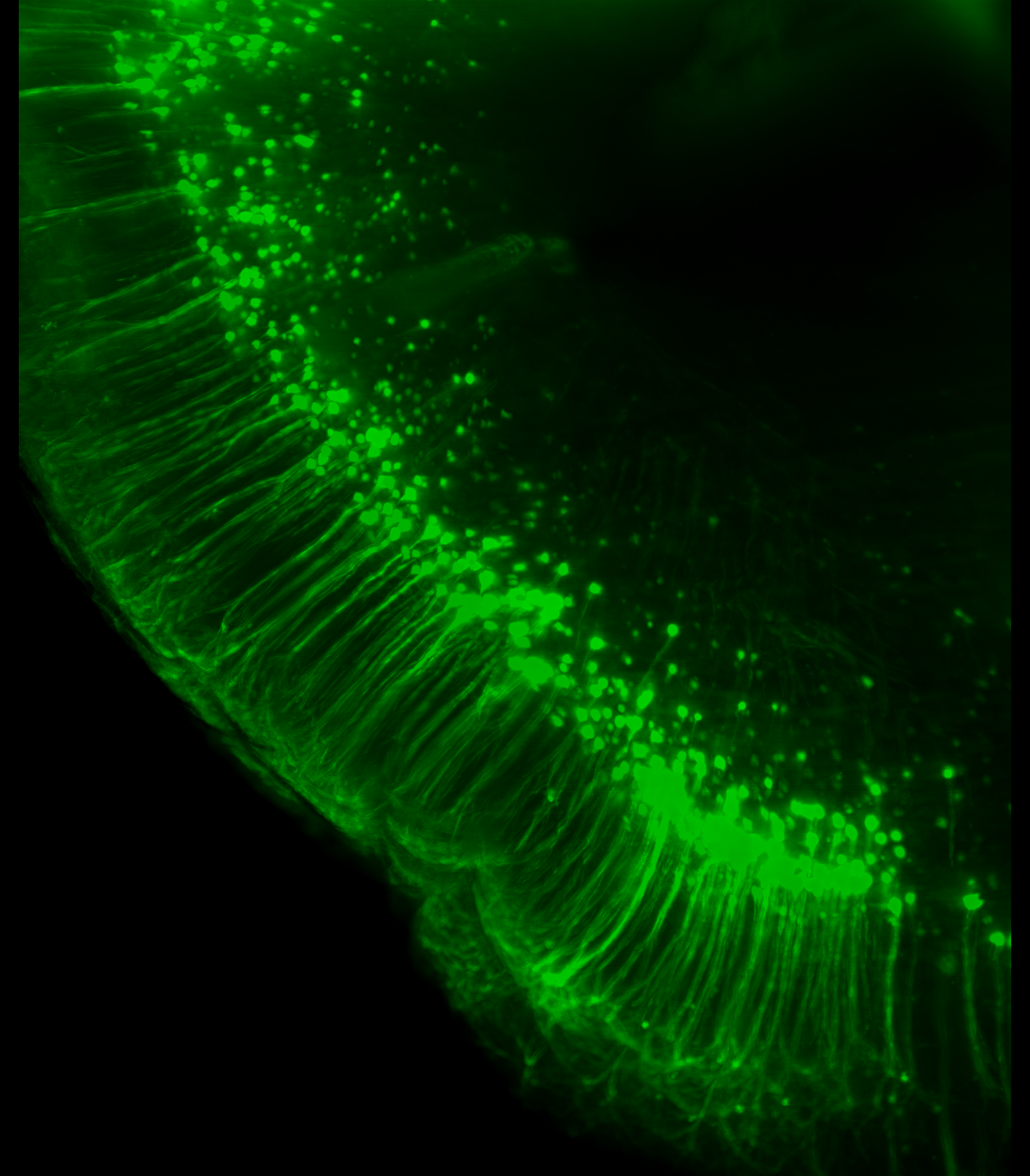
